# Senescent human fibroblasts have increased FasL expression and impair the tumor immune response

**DOI:** 10.1101/2024.06.10.598270

**Authors:** Monica Cruz, Joshua Dulong, Georgio Mansour Nehmo, Anthony Sonn, Gaël Moquin-Beaudry, Basma Benabdallah, Oanh Le, Christian Beauséjour

## Abstract

Syngeneic mouse tumor models have shown that senescence influences the tumor immune response in multiple ways, including the induction of an immunosuppressive microenvironment or the promotion of immune cell recruitment. Yet, the impact of senescence on the tumor immune response in a humanized setting remains largely unexplored. To address this, we employed tumor spheroids and mice bearing tumors immunogenic to human immune cells derived from the same donor. We found that senescent fibroblasts exert a dual effect by enhancing the recruitment of immune cells into the tumor microenvironment while simultaneously promoting the apoptosis of T and NK cells. Mechanistically, we demonstrate that this apoptosis is primarily due to increased Fas ligand (FasL) expression on the surface of senescent fibroblasts. Deletion of FasL on fibroblasts was sufficient to prevent immune cell death and increase tumor cell killing. Our results highlight the importance of evaluating the impact of therapy-induced senescence in humanized models to understand and predict the outcome of cancer treatments.

## INTRODUCTION

Cancer treatments can induce senescence of cancer cells and other cell types within the tumor microenvironment (TME) (1). This phenotype is believed to have mostly detrimental consequences, as senescent cells were shown to play a significant role in the development and progression of cancer (2, 3). Indeed, Krtolica and colleagues were the first to demonstrate that senescent human fibroblasts favor tumor growth in immune-deficient mice (2). The accumulation of senescent fibroblasts was also shown to contribute to local and systemic inflammation, promoting adverse effects and cancer relapse (4). These effects are believed to be mediated mainly by the senescence-associated secretion phenotype (SASP), which includes several factors linked to inflammation and malignancy (5, 6).

Several studies using syngeneic tumor models in immune-competent mice showed that the SASP can prevent tumor rejection by inducing an immunosuppressive TME (7–10). For example, prolonged exposure to SASP factors such as type 1 interferon and IL-6 were shown to downregulate immune cell functions (11–13) and to interfere with the tumor immune response mediated by T and NK cells (7–9). On the other hand, other studies showed that the SASP can stimulate and recruit immune cells (14–17). Hence, while the SASP can undoubtedly promote tumor growth, its impact on immune cells is variable and likely dependent on the tumor model.

Several immune cell populations impact tumor growth, including cytotoxic T cells, whose intra-tumoral presence correlates with a survival benefit (18). However, the tumor immune response is known to be significantly diminished by myeloid-derived suppressor cells (MDSC) and other immune-inhibitory cells or molecules within the TME (19, 20). One of these factors is FasL, which can trigger apoptosis of Fas-expressing effector cells such as T cells and natural killer (NK) cells (21). FasL is expressed by several cell types, including tumor cells, MDSC (20, 21), tumor-associated macrophages, regulatory T cells (21), and cancer-associated fibroblasts (22). Elevated FasL levels in solid tumors correlate with disease progression, increased metastasis, poor overall survival (23), and decreased numbers of CD8^+^ infiltrating T cells (24).

Nevertheless, the impact of senescence on the tumor immune response in a humanized setting remains undetermined. This is because, until recently, there was no adequate tumor model where human cancer cells could be rejected or their growth delayed in the presence of immune cells. Such models are essential to evaluate whether senescent cells interfere with the tumor immune response and if so, through which mechanism. To this end, we recently developed iPSC-derived humanized mouse models where tumors can be rejected following the injection of autologous human immune cells (25). Using these models, we show here that senescent human dermal fibroblasts (HDF) have a dual impact on the tumor immune response. On the one hand, senescent HDF attract immune cells; on the other hand, they induce their death. Mechanistically, we demonstrate that this is mostly the result of increased FasL expression at the surface of senescent HDF. By providing a better understanding of the complex interactions between senescent stromal cells and immune cells, our results should help develop better-targeted therapeutic interventions.

## RESULTS

### Senescent fibroblasts promote tumor growth and stimulate the infiltration of immune cells in humanized mice

To determine the role of a senescent stroma on the tumor-immune response, we first induced senescence of HDF by exposure to ionizing radiation (IR) or the expression of K-RAS^V12^, two proven methods that we and others routinely use (**Supplementary Fig. 1A and B)** (26, 27). We then injected non-senescent or senescent HDF together with tumor cells into the flanks of NSG-SGM3 mice (ratio of 1:4). We used two tumor cell lines (HEPA-4T and LEC-4T derived from hepatic and lung iPSC-derived progenitor cells respectively) which we previously showed are immunogenic to autologous immune cells (25). The day after tumor inoculation, mice were injected intraperitoneally with human immune cells (PBMCs and granulocytes, 5×10^6^ of each). Note that tumor cells, HDF, and immune cells were all derived from the same donor to evaluate the impact of senescence in an autologous setting (**Fig. 1A**). HEPA-4T and LEC-4T tumor cells were transduced to express mPlum and HDF were stained with the NIR790 cytoplasmic membrane dye, allowing the visualization of both cell types, and tracking of tumor growth by quantitative *in vivo* imaging (**Fig. 1B**).

**Figure 1.**
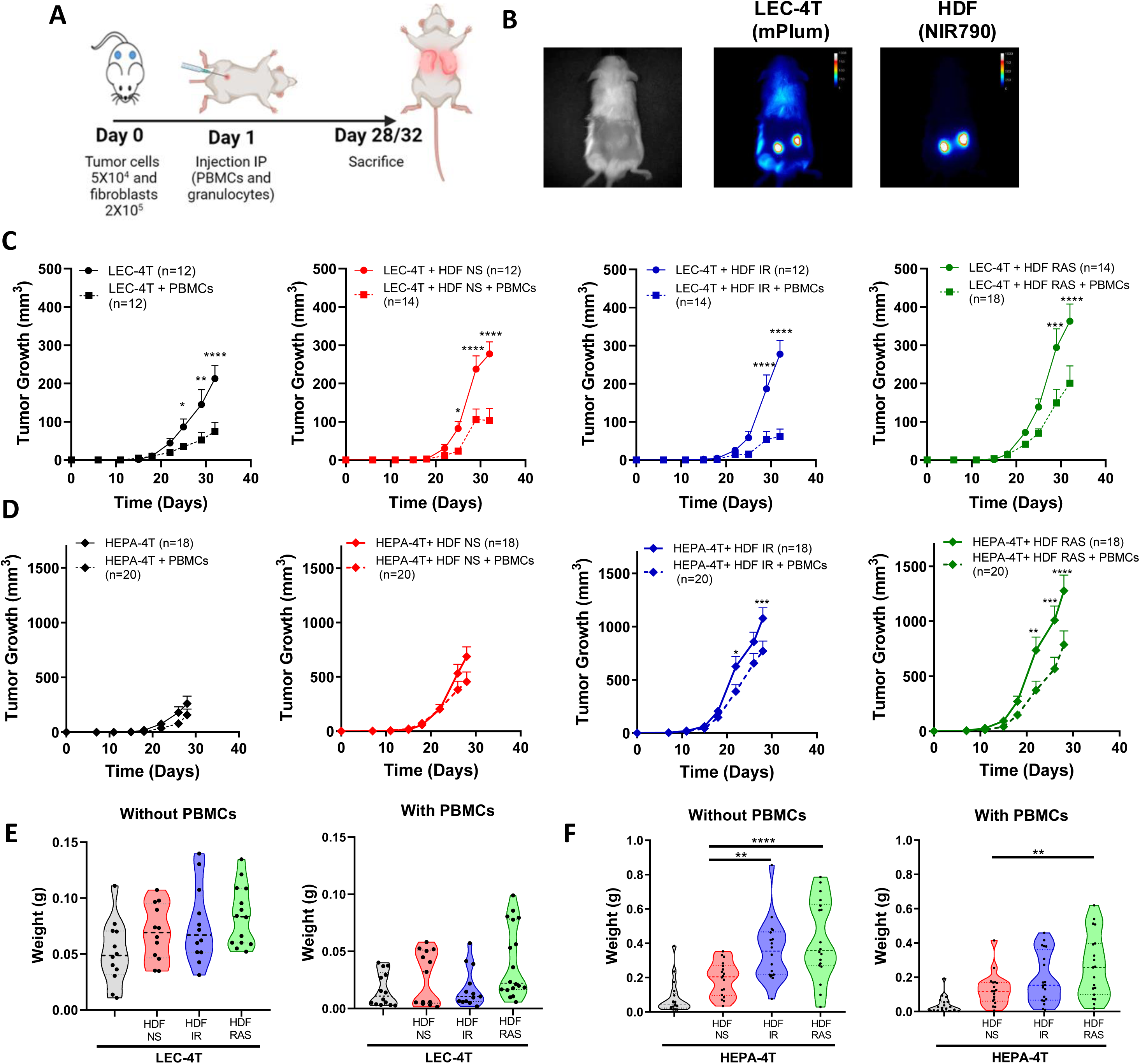
Senescent fibroblasts promote tumor growth without compromising the tumor immune response in humanized mice. **A)** Schematic of the *in vivo* experimental design. NSG-SGM3 mice were subcutaneously injected with LEC-4T or HEPA-4T tumor cells (5 × 10^4^) and non-senescent or senescent HDF (2 × 10^5^ cells) on day 0. The next day, mice were injected intraperitoneally with PBMCs (5 × 10^6^) and granulocytes (5 × 10^6^). Mice were sacrificed on day 28 or 32 depending on tumor cells injected and tumors were collected for analysis. **B)** Representative images of LEC-4T tumors on day 22 expressing mPlum and HDF stained with the NIR790 dye after their injection. **C)** Growth curves for LEC-4T tumors injected alone (in black n=12) or co-injected with non-senescent HDF (in red n=12-14), senescent HDF induced by irradiation (in blue n=12-14) or induced by RAS (in green n=14-18) in mice without (solid line) and with autologous immune cells (dashed line). Each line represents the mean tumor growth (± SEM) over 32 days. Statistical analyses were performed using a mixed-effects model, followed by Tukey’s multiple comparison test. **D)** Growth curves for HEPA-4T tumors injected alone (in black n=18-20) or co-injected with non-senescent HDF (in red n=18-20), senescent HDF induced by irradiation (in blue n=18-20) or induced by RAS (in green n=18-20) in mice without (solid line) and with autologous immune cells (dashed line). Each line represents the mean tumor growth (±SEM) over 28 days. Statistical analyses were performed using a mixed-effects model, followed by Tukey’s multiple comparison test. **E and F)** Graphs representing the weights of LEC-4T or HEPA-4T tumors collected at sacrifice. Each dot represents an individual tumor. Values represent the mean ± SEM. A one-way ANOVA with Dunnett’s multiple comparisons tests was used to determine statistical significance.

As expected, the co-injection of tumor cells with non-senescent or senescent HDF resulted in accelerated tumor growth compared to mice injected with tumor cells alone (**Fig. 1C-D**). The injection of immune cells was able to delay tumor growth in all groups. Surprisingly, the presence of HDF did not seem to interfere with the tumor rejection, except for RAS-induced HDF, which slightly hampered the rejection of LEC-4T tumors. Indeed, for this group, tumor size was reduced only by about 45% in the presence of immune cells compared to 60-80% in the other groups (**Fig. 1C**). These results were confirmed at the time of sacrifice when residual tumors were excised and weighted (**Fig. 1E and 1F**). *In vivo*, imaging of NIR790-labeled HDF over four weeks following the injection of immune cells revealed that most HDF persists over time, suggesting they are not the target of immune cells (**Supplementary Fig. 1C and D**). Overall, these results demonstrate that senescent HDF have either no effect or only a moderate effect on the tumor immune rejection in these humanized models.

We anticipated that senescent HDF, through a combination of their mitogenic effect and immunosuppressive features, would interfere with tumor rejection. Hence, to better understand the impact of senescence, we characterized the infiltration of human immune cells (hCD45^+^) in dissociated tumors at the time of sacrifice. Cell counts obtained by flow cytometry and normalized to the tumor weight showed that tumors injected with HDF, particularly senescent HDF, tended to have an increased infiltration of hCD45^+^ immune cells. Specifically, LEC-4T injected with IR-induced senescent HDF showed a twofold increase in immune cell infiltration, while LEC-4T or HEPA-4T injected with RAS-induced senescent HDF demonstrated a fourfold and twofold increase, respectively, compared to tumors injected with non-senescent HDF (**Fig. 2A and 2C**). However, while the ratio of tumor-infiltrating immune cell types was slightly different between the two tumor cell lines, it was not significantly changed between groups and was composed mainly of CD4^+^ and CD8^+^ T cells (**Fig. 2B and 2D**). Because residual tumors had different sizes at the time of sacrifice, which likely impacted infiltration, we then measured immune cell infiltration using a more controlled spheroid model (**Fig. 2E**). Therefore, LEC-4T and HEPA-4T cell lines expressing mPlum were cultured in 3D spheres with or without non-senescent or senescent HDF expressing GFP (**Fig. 2F**). Spheroids were co-cultured for 24 hours with PBMCs, after which the localization of infiltrated immune cells into the spheroids was assessed by immunofluorescence. Images showed an increase in the number of infiltrating immune cells, particularly towards the core of the spheroids, in the presence of senescent HDF (**Fig. 2G**). To better quantify the infiltrated cells, we repeated the experiment, but this time, we dissociated the spheres with trypsin and counted cells by flow cytometry. Increased immune cell infiltration in the presence of senescent HDF was confirmed (**Fig. 2H**). These observations led us to hypothesize that senescent HDF may have opposing effects as they can induce the recruitment of immune cells yet without enhancing tumor rejection.

**Figure 2.**
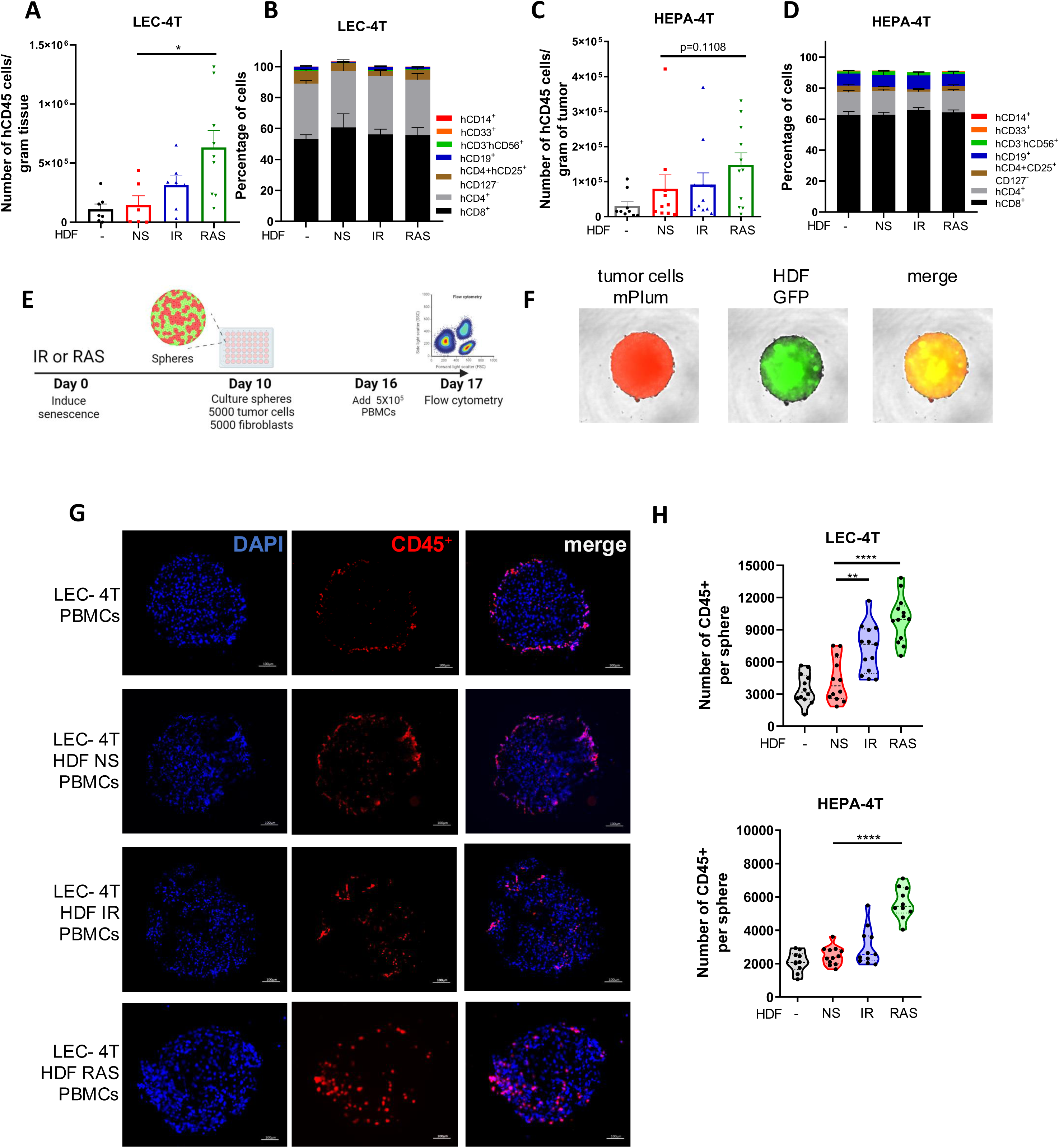
Senescent fibroblasts increase immune cell infiltration in spheroids and in tumors. **A)** Graph showing the absolute counts of tumor-infiltrating human CD45^+^ cells per gram of dissociated LEC-4T tumors as determined by flow cytometry. Each dot represents the count from a single tumor. Shown is the mean ± SEM. Statistical analysis between groups was performed with one-way ANOVA followed by Dunnett’s multiple comparisons tests. **B)** Stacked bar graph showing the proportion of immune cell subset populations in LEC-4T tumors (the same as in panel A) from each group. Shown is the mean ± SEM. **C)** Graph showing the absolute counts of tumor-infiltrating human CD45^+^ cells per gram of dissociated HEPA-4T tumors as determined by flow cytometry. Each dot represents the count from a single tumor. Shown is the mean ± SEM. Statistical analysis between groups was performed with one-way ANOVA Dunnett’s multiple comparisons tests. **D)** Stacked bar graph showing the proportion of immune subset cell populations in HEPA-4T tumors (the same as in panel C) from each group. Shown is the mean ± SEM. **E)** Schematic representation of the 3D tumor spheroid invasion model. In brief, HDF were exposed or not to IR or RAS and 10 days later, an equal number of tumor cells (LEC-4T or HEPA-4T) were mixed with non-senescent or senescent HDF (5000 cells each) to form mixed-cell spheroids. Once the spheroid assembled and matured for 6 days, 5 × 10^5^ human PBMCs were added and the infiltration of immune cells into the spheroid was quantified 24 hours later by flow cytometry. **F)** Representative images (at 4X) of spheroids at day 16 before adding immune cells. Tumor cells are shown in red (mPlum), HDF in green (GFP) and in yellow the merge of the two signals. **G)** Representative images of immunostained sections from the indicated spheroids showing infiltrated human immune cells (CD45^+^ in red) and nuclei (DAPI in blue). The scale bar represents 100 μm. **H)** Graphs showing the number of human immune cells (CD45^+^) infiltrated in spheroids as determined by flow cytometry from each indicated group. Each dot represents the count from a single dissociated sphere obtained in three independent experiments. Statistical analysis between groups was performed with one-way ANOVA with Dunnett’s multiple comparisons tests.

### Senescent fibroblasts trigger apoptosis of human immune cells through increased expression of FasL

To better evaluate the effects of senescent HDF on immune cells, we used a model where PBMCs collected from three healthy donors were co-cultured for up to 72 hours with non-senescent or senescent HDF. Immune cells were then counted either by flow cytometry or alternatively, immune cell death was assessed with propidium iodide (PI) staining and quantified using live-cell imaging (**Fig. 3A**). For all donors, we found a substantial decrease in the absolute number of live CD45^+^ in contact with senescent HDF, mostly RAS-induced senescent HDF (**Fig. 3B**). Such a decrease in cell viability was primarily due to dying CD3^+^ T cells and to a lesser extent CD3^−^CD56^+^ NK cells although the effect was donor dependant (**Fig. 3B**). This ability of senescent HDF to kill immune cells was also confirmed by counting dying (PI positive) immune cells using live-cell imaging (**Fig. 3C and 3D**).

**Figure 3.**
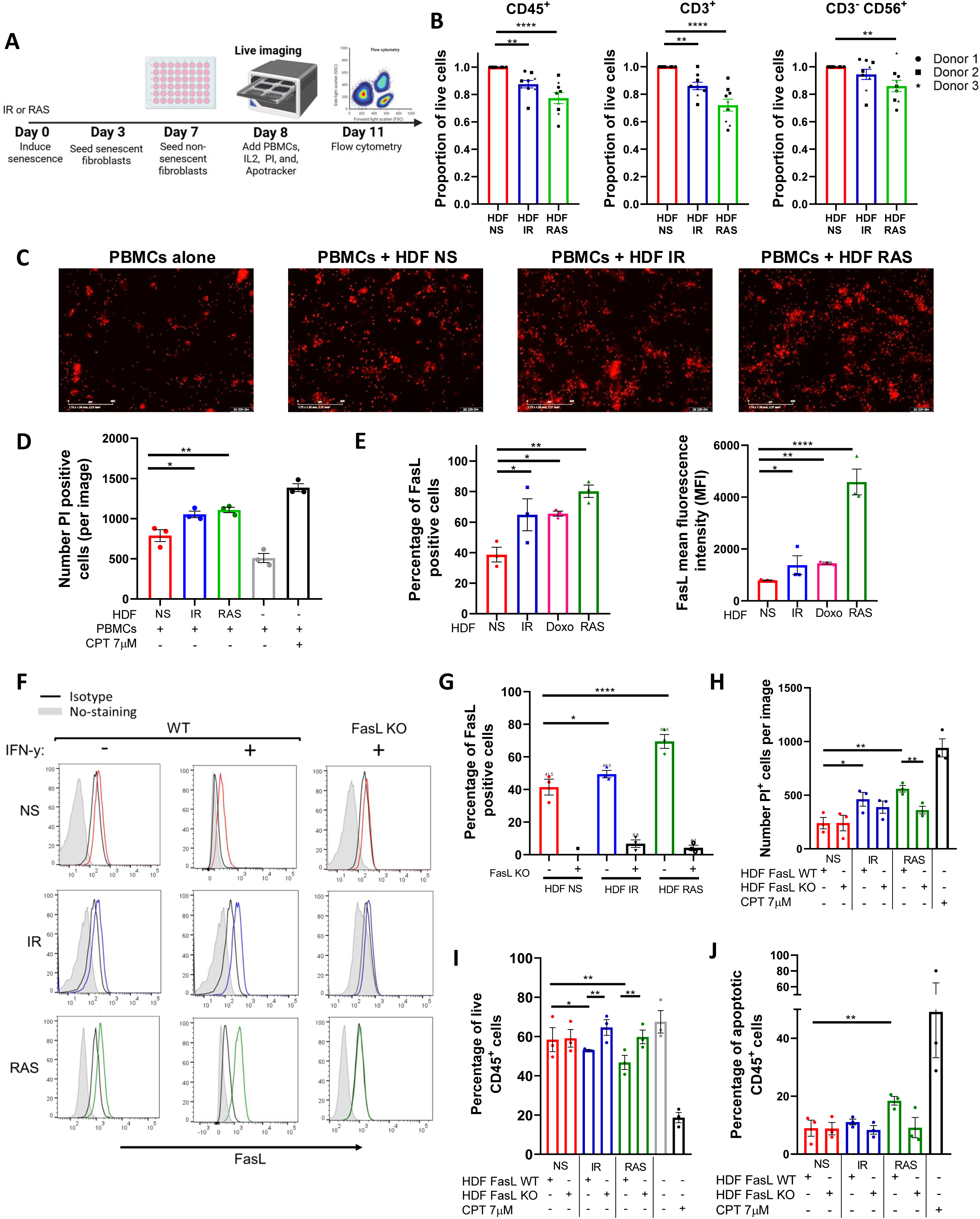
Senescent fibroblasts induce immune cell death through up-regulation of FasL. **A)** Schematic representation of the experimental design. In brief, HDF were exposed to IR or RAS, and 3 days later, 3 × 10^4^ cells were seeded in a 48-well plate. An equal number of non-senescent HDF were seeded only on day 7, and the day after PBMCs (3 × 10^5^) were added. Cell death as determined by propidium iodide (PI) incorporation tracked over 72 hours at 2-hour intervals using an IncuCyte live-cell analysis system or alternatively, absolute cell counts were determined by flow cytometry at 72 hours. **B)** Graphs showing the proportion of live PBMCs (CD45^+^), T cells (CD3^+^), and NK cells (CD3^−^CD56^+^) after 72 hours of coculture with senescent HDF normalized to counts obtained in the presence of non-senescent HDF. Data are presented as mean ± SEM from three independent experiments conducted with each donor of PBMCs, utilizing three distinct donors. Statistical analysis between groups was performed, followed by one-way ANOVA with Dunnett’s multiple comparisons tests. **C)** Representative images captured by the IncuCyte using a 10X objective showing PI^+^ (in red) dying cells at 72 hours. The scale bar represents 400 µm. **D)** Quantification of PI^+^ dying immune cells from each group. CPT was added as a positive control. Data are presented as mean ± SEM from three independent experiments. Statistical analysis between groups was performed with one-way ANOVA followed by Dunnett’s multiple comparisons tests. **E)** Quantification by flow cytometry of the proportion and the mean fluorescence intensity (MFI) of non-senescent and senescent HDF expressing FasL at their surface. The MFI was calculated by subtracting the fluorescence of the isotype control. Data are presented as mean ± SEM from three independent experiments. Statistical analysis between groups was performed with one-way ANOVA with Dunnett’s multiple comparisons tests. **F)** FasL expression as detected by flow cytometry on wild type (WT) and FasL KO HDF either non-senescent (NS in red) or senescent (IR in blue or RAS in green). Also shown are non-stained cells (in gray) and cells stained with the isotype controls (in black). Cells were stimulated or not with IFN-γ (200ng/mL). Representative histograms of three independent experiments are shown. **G)** Bar graphs showing the proportion of cells expressing FasL in the different HDF populations presented in panel F. Data are presented as mean ± SEM from three independent experiments. Statistical analysis between groups was performed with one-way ANOVA with Dunnett’s multiple comparisons tests. **H)** Quantification of PI^+^ dying immune cells after 72 hours of co-culture with the indicated HDF populations. Data was acquired with the IncuCyte® live-cell analysis system. CPT was added as a positive control. Data are presented as mean ± SEM from three independent experiments. The p-value was calculated by multiple t-tests. **I)** Quantification of live (Annexin V^−^/PI^−^) CD45^+^ cells after 72 hours of co-culture with the indicated HDF populations as determined by flow cytometry. CPT was used as a positive control. Data are presented as mean ± SEM from three independent experiments. Statistical analysis was calculated by multiple t-tests. **J)** Quantification of apoptotic (Annexin V^+^/PI^+^) CD45^+^ cells after 72 hours of co-culture with the indicated HDF populations as determined by flow cytometry. CPT was used as a positive control. Data are presented as mean ± SEM from three independent experiments. Statistical analysis was calculated by multiple t-tests

We next wanted to investigate the mechanism leading to immune cell death. We found that senescent HDF consistently increased FasL expression independently of the inducer (**Fig. 3E**). Such an increase in the proportion of cells expressing FasL was also accompanied by a significant two and five-fold increase in the mean fluorescence intensity in IR/doxorubicin and RAS-induced senescent HDF respectively (**Fig. 3E**). Similar results were observed using other human fibroblast cell lines (**Supplementary Fig. 2**). To confirm the involvement of FasL in immune cell killing, we generated FasL knockout (KO) HDF using CRISPR-Cas9 and confirmed loss of FasL expression by flow cytometry in the absence or presence of IFN-γ which increases FasL expression in wild-type cells (**Fig. 3F and 3G**). We then performed co-culture studies using HDF and human PBMCs, and we observed that immune cells survival was not compromised in contact with FasL KO HDF as determined by counting the number of PI+ cells per image using the IncuCyte software (**Fig. 3H**). Increased immune cell survival in contact with FasL KO senescent HDF was also confirmed using flow cytometry (**Fig. 3I**). Staining for Annexin V showed that immune cell death was induced mainly by apoptosis, especially by RAS-induced HDF (**Fig. 3J**). Overall, these findings indicate that immune cells killing by senescent HDF, particularly CD3^+^ T cells and CD3^−^CD56^+^ NK cells, is mostly mediated by increased FasL expression.

### Therapy-induced senescence increases FasL expression and decreases immune cell survival

We next wanted to evaluate if increased FasL expression would occur in the context of therapy-induced senescence and if this would negatively impact immune cell survival. Therefore, we generated tumor spheroids by mixing an equal number of HDF and A549 tumor cells (**Fig. 4A**). We chose to work with the A549 lung epithelial carcinoma cell line for this experiment because A549 cells adopt a senescence phenotype when exposed to IR or doxorubicin. Indeed, we observed that A549 spheroids, after treatment, increased SA-β-gal activity and significantly reduced their growth (**Fig. 4B and C**). Immunofluorescence imaging performed on cryosections showed an increase in FasL expression in response to treatments in tumor spheroids (**Fig. 4D**). Considering the relatively low abundance of HDF in our spheres on day 11 (day 6 after treatment), and nearly no increase in FasL in treated A549 spheroids in absence of immune cells (**Fig. 4D and Supplementary Fig. 3**), the observed increase in FasL expression was higher than expected. Such an increase may be attributed to the 3D environment or derive from infiltrating immune cells. Nonetheless, when we analyzed the proportion of live immune cells (CD45^+^, PI^−^) in dissociated spheroids we observed significantly less viable immune cells in the context of therapy-induced senescence (**Fig. 4E**). To determine whether FasL expression is also upregulated in vivo, we injected HDF intravenously into mice and allowed them to engraft in the lungs, the primary site of localization following injection, before subjecting mice to irradiation. (**Fig. 4F**). This model allowed us to easily retrieve the injected HDF in mice. As observed *in vitro*, lung tissue sections containing HDF had increased FasL expression compared to sections collected from non-irradiated mice (**Fig. 4G-H**).

**Figure 4.**
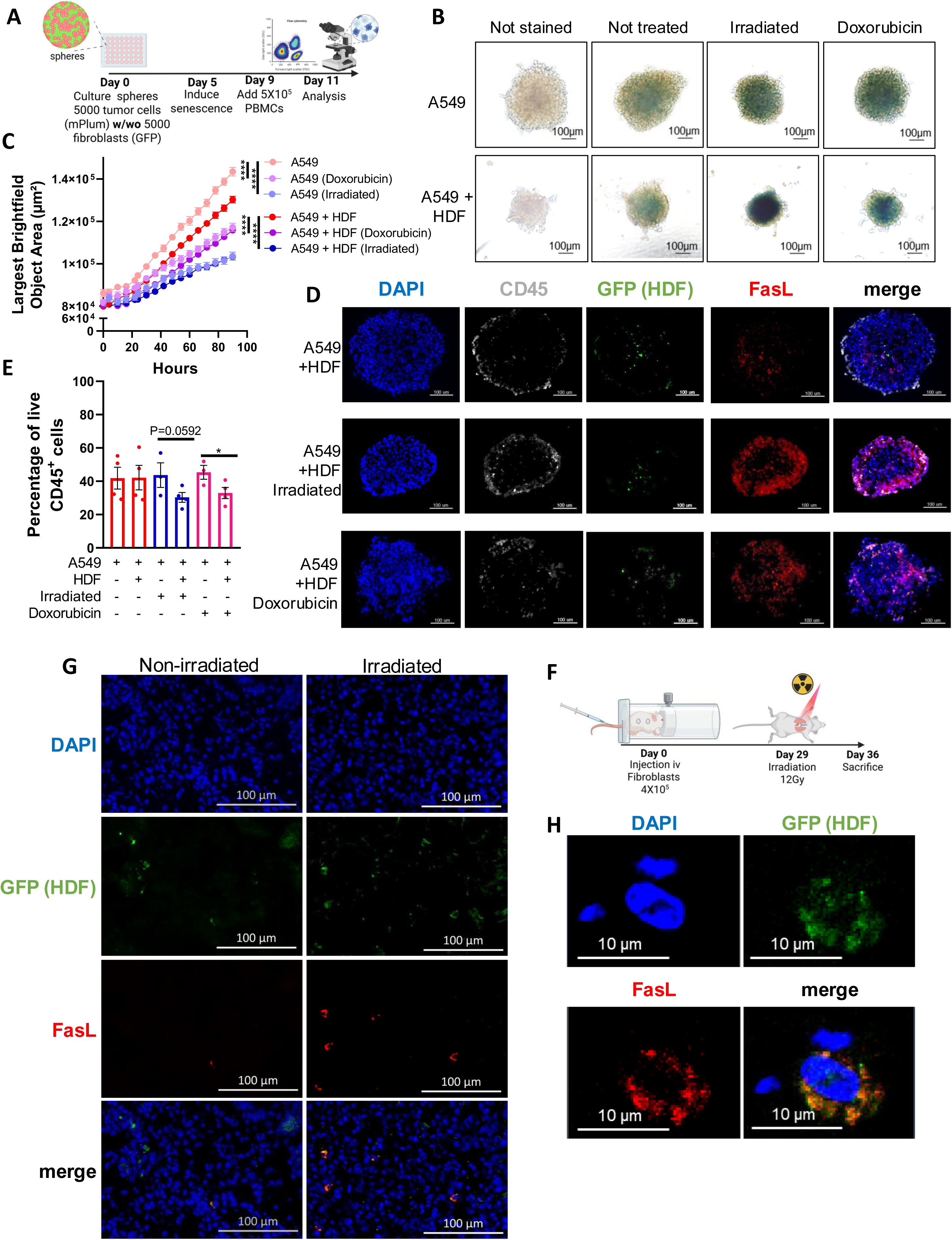
Therapy-induced senescence in tumor spheroids impairs the viability of infiltrated immune cells. **A)** Schematic representation of the 3D tumor spheroid model. In brief, A549 tumor cells alone or with an equal number of HDF (5000 cells for each) were mixed to form monospheroids or mixed-cell spheroids, respectively. Five days after, spheroids were treated with doxorubicin (0.1µM) or irradiated (15 Gy) to induce senescence. Then, four days later PBMCs (5 × 10^5^) were added, and their infiltration into the spheroid was quantified 48 hours later by flow cytometry. **B)** Representative images of spheroids stained for β-galactosidase activity six days after being treated with doxorubicin or irradiation. Unstained spheroids or spheroids not exposed to therapy were used as controls. The scale bar represents 100 μm. **C)** Graph showing the average size of spheroids represented by the largest brightfield object area metric (µm^2^) over time following treatments as detected by IncuCyte imaging. Shown is the mean ± SEM of three independent experiments. Statistical differences were identified by mixed-effects modeling with Tukey’s multiple comparisons. **D)** Representative images of immunostained spheroid sections showing CD45⁺ immune cell infiltration (white), the presence of HDFs (GFP, green), FasL expression (red), and cell nuclei (DAPI, blue). Scale bar = 100 μm. **E)** Graph showing the proportion of live (Annexin V^−^/PI^−^) CD45^+^ immune cells infiltrated in spheroids from the indicated groups. Each dot represents the average of infiltrated cells in n=6 spheroids collected from four independent experiments. Shown is the mean ± SEM. Statistical analysis between groups was performed by a one-way ANOVA with Tukey’s multiple comparisons. **F)** Schematic of the in *vivo* experimental design. NSG-SGM3 mice were injected intravenously (i.v) with 4X10^5^ non-senescent HDF. After 29 days mice received a single dose of whole thorax radiation (12Gy). Seven days after, mice were sacrificed, and lungs were collected for analysis. **G)** Representative immunofluorescence staining of HDF (GFP in green), FasL (in red), and cell nuclei (DAPI in blue) from irradiated and non-irradiated lung tissues. The scale bar represents 100 μm. **H)** Confocal images shown at higher magnification of a tissue section as described in panel G.

### The SASP suppresses tumor immunity through FasL-dependent and independent mechanisms

The SASP allows for a crosstalk between senescent cells and neighboring cells. Therefore, we next wanted to determine if the SASP alone could induce immune cell death, knowing that FasL can be found soluble (28, 29). We observed that soluble FasL was indeed present in the conditioned media (CM) of RAS-induced senescent HDF (63 pg/mL) but not in any other conditions, including in CM from IR-induced senescent HDF (**Fig. 5A**). Therefore, to determine the impact of the SASP to induce death cell, we incubated immune cells in the presence of CM collected from non-senescent and senescent HDF for 48 hours and counted the number of viable immune cells (CD45^+^ and CD3^+^) by flow cytometry. In accordance with the results of the ELISA, we found a significant decrease in the number of viable immune cells in the presence of CM collected from RAS-induced senescence HDF (**Fig. 5B**). Nonetheless, we noted a modest reduction in cell survival when exposed to conditioned media from IR-induced senescent HDF. This indicates that the SASP has a toxic effect on immune cells, independent of soluble FasL, albeit to a lesser degree (**Fig. 5B**).

**Figure 5.**
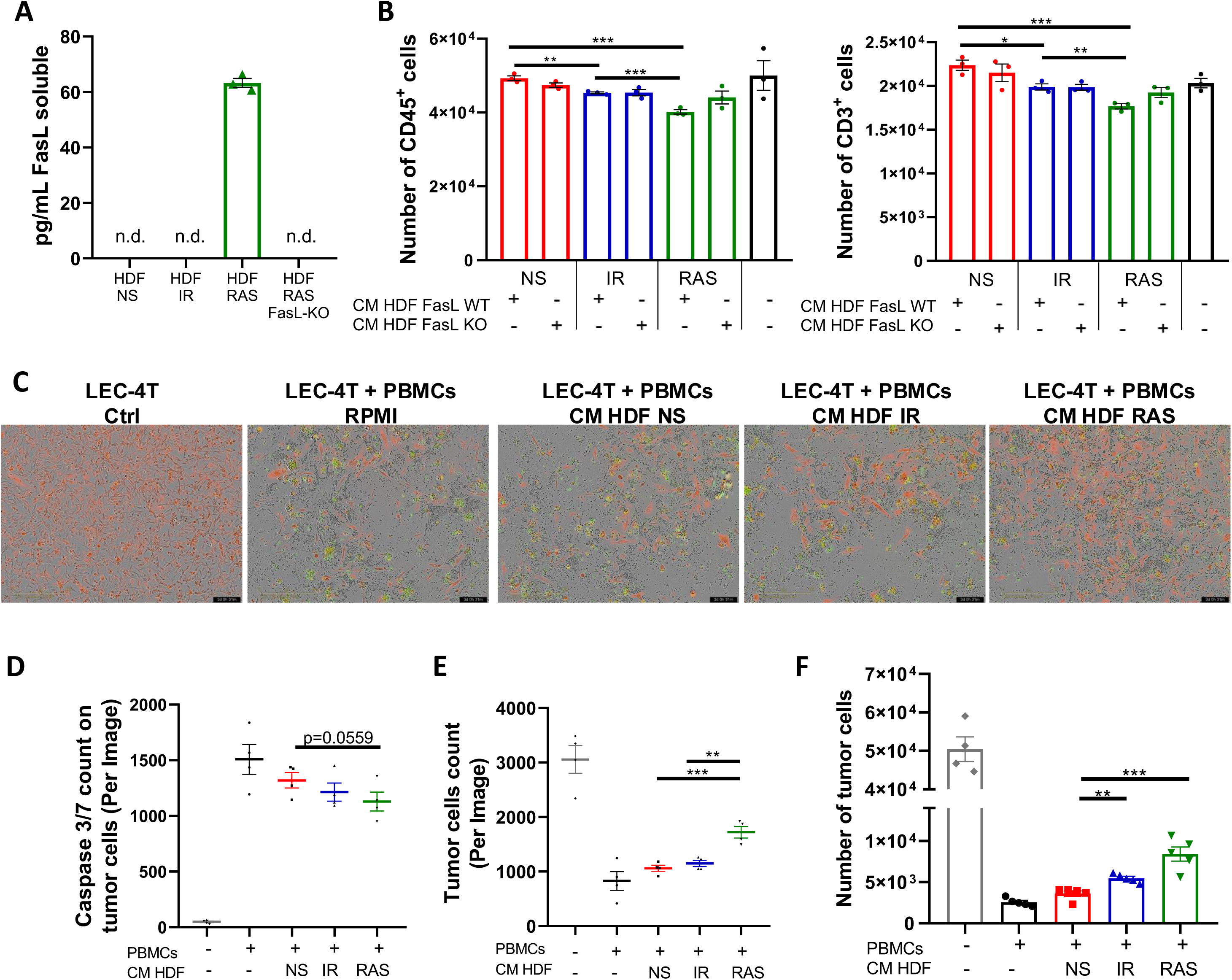
The SASP triggers apoptosis of immune cells and promotes tumor cell survival *in vitro*. **A)** Concentration of soluble FasL (sFasL) as detected by ELISA in the conditioned media (CM) of non-senescent and senescent HDF populations. Each dot represents an independent experiment (n=3), and t-tests determined statistical significance between groups. **B)** Graphs showing the absolute counts of CD45^+^ (left panel) and CD3^+^ (right panel) live cells as determined by flow cytometry after 48 hours of culture in CM collected from the indicated HDF populations. Shown is the mean ± SEM from three independent experiments. Multiple t-tests determined statistical significance between groups. **C)** Representative images of LEC-4T tumor cells (expressing mPlum in red) after 72 hours co-cultured with PBMCs in CM collected from the indicated HDF populations. The Caspase-3/7 Green reagent was added to detect cells undergoing apoptosis. Scale bars represent 400 μm. **D)** Scatter dot plot showing the number of caspase 3/7 positive tumor cells (red and green overlap) after 72 hours of co-culture as determined using the IncuCyte® live-cell analysis software. Shown is the mean ± SEM from four independent experiments. Multiple t-tests determined statistical significance between groups. **E)** Scatter dot plot showing the number of LEC-4T tumor cells (in red) after 48 hours of co-culture as determined using the IncuCyte® live-cell analysis software. Shown is the mean ± SEM from four independent experiments. Multiple t-tests determined statistical significance between groups. **F)** Absolute counts of live LEC-4T cells as measured by flow cytometry at the end of the 72-hours co-culture period. Each dot represents the average of five technical replicates. Shown is the mean ± SEM from four independent experiments. Multiple t-tests determined statistical significance between groups.

We also observed that the SASP had immunosuppressive properties as it prevented the proliferation of CFSE-labelled immune cells in a mixed lymphocyte reaction. For this experiment, PBMCs from two unrelated donors were mixed in the presence or absence of CM collected from non-senescent or senescent HDF, and the proliferation of T cells was analyzed by flow cytometry. The results showed that the SASP can interfere with the proliferation of T cells (**Supplementary Fig. 4**). Considering these observations, we reasoned that increased immune cell killing combined with their reduced proliferation should lead to reduced killing of tumor cells. To test this hypothesis, we incubated PBMCs with LEC-4T tumor cells (expressing mPlum) in the absence or presence of CM for 72 hours. Using live-cell imaging, we tracked tumor cell growth and apoptosis (Caspase 3/7 specific dye in green). As expected, images showed less apoptosis of tumor cells and a consequent increase in their number in the presence of CM collected from senescent HDF (**Fig. 5C-E**). In agreement with soluble FasL being present only in CM collected from RAS-induced senescent HDF, tumor cell counts were higher in this group compared to IR-induced senescent HDF. Counting live tumor cells by flow cytometry confirmed the results (**Fig. 5F**). In line with these observations, the killing of tumor cells by primary human NK cells was also found inhibited when placed in co-culture with wild type but not FasL KO HDF (**Supplementary Fig. 5**). Together, this demonstrates that senescent HDF, partly through their SASP, can protect tumor cells either by killing immune cells or by limiting their proliferation using both FasL-dependent and independent mechanisms.

### FasL KO senescent fibroblasts do not interfere with the tumor immune response in humanized mice

Finally, we evaluated if tumors would be better rejected by immune cells in the absence of FasL expression from senescent HDF. To do so, we used LEC-4T tumor cells co-injected with RAS-induced senescent HDF, the combination with the highest impaired tumor-immune rejection (**Fig. 1C**). We co-injected subcutaneously LEC-4T cells with either WT or FasL KO RAS-induced senescent HDF in NSG-SGM3 mice (**Fig. 6A**). First, we confirmed that in the absence of immune cells, that WT or FasL KO senescent HDF had a similar mitogenic effect on tumor growth when compared to tumor cells injected alone (**Fig. 6B and 6C**). However, in the presence of immune cells, we observed that tumors containing FasL KO senescent HDF were better rejected, to a level similar to the one observed in tumors without HDF (**Fig. 6B and 6C**). Tumor weight at the time of sacrifice confirmed these results (**Fig. 6D**). However, the number and proportion of tumor-infiltrating immune cells were surprisingly not significantly changed in the absence of FasL expression (**Fig. 6E and 6F**). We speculate that this may be the consequence of the relatively small size of residual tumors in the FasL KO group at the time of sacrifice, which may not accurately represent the peak of immune cell infiltration during the tumor immune response.

**Figure 6.**
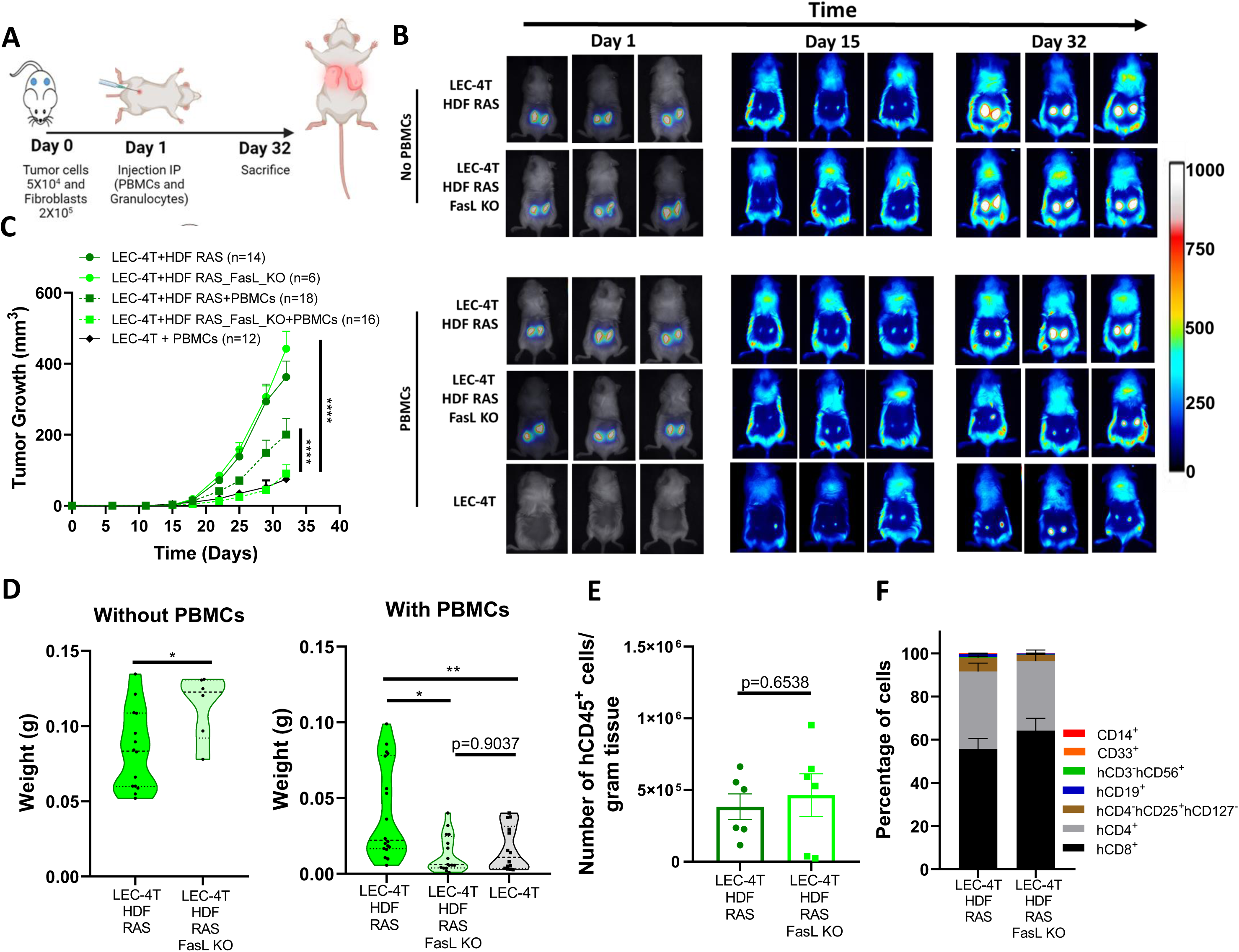
FasL KO senescent fibroblasts do not disrupt the tumor immune response in humanized mice. **A)** Schematic of the in *vivo* experimental design. NSG-SGM3 mice were subcutaneously injected with LEC-4T tumor cells (5 × 10^4^ ) and non-senescent or senescent HDF (2 × 10^5^ cells) on day 0. The next day, mice were injected intraperitoneally with PBMCs (5 × 10^6^) and granulocytes (5 × 10^6^). Mice were sacrificed on day 32 and tumors were collected for analysis. **B)** Representative images of mice bearing subcutaneous LEC-4T tumors (expressing mPlum) alone or co-injected with the indicated HDF populations stained with the NIR790 dye after their injection. Images from day 1 show HDF-stained with the NIR790 dye while days 15 and 32 show mPlum tumor growth. **C)** Growth curves for LEC-4T tumors in humanized mice injected with LEC-4T cells alone (n=12) or co-injected with RAS-induced senescent HDF (n=18) or RAS-induced senescent HDF KO for Fas-L (n=16). Also shown is the growth curve of LEC-4T tumors in NSG-SGM3 mice co-injected with RAS-induced senescent HDF (n=14) or RAS-induced senescent HDF KO for Fas-L (n=6). Each line represents the mean tumor growth (±SEM) over 32 days. Statistical analyses were performed using a mixed-effects model, followed by Tukey’s multiple comparison test. **D)** Shown is the weight of LEC-4T tumors collected at sacrifice on day 32. Each dot represents the weight of an individual tumor. Values represent the mean ± SEM. Statistical analysis between groups was performed by t-test or one-way ANOVA with Tukey’s multiple comparisons. **E)** Bar graph showing the number of tumor-infiltrating hCD45^+^ immune cells per gram of tumor collected from both groups, RAS-induced senescent HDF WT and FASL KO, at the time of sacrifice. Each dot represents the infiltration in an individual residual tumor large enough to be excised and dissociated (n=6). Values represent the mean ± SEM, and a t-test was performed for statistical analysis. **F)** Stacked bar graph showing the proportion of immune cell subset populations in LEC-4T tumors collected from the indicated groups. Shown is the mean ± SEM.

## DISCUSSION

Using immunogenic autologous tumors, we were able to uncover a dual role of senescence on the tumor immune response in humanized mice. Indeed, we observed that senescent HDF, aside from their mitogenic effect on tumor cells, can recruit immune cells and then induce their death through a mechanism that involves FasL. Importantly, we observed increased FasL expression on various HDF lines in response to different senescence inducers. Highlighting the importance of the tumor model to study the impact of senescence on the tumor immune response, we found that subcutaneous A549 tumors were not rejected despite being infiltrated by immune cells at similar levels to those observed in HEPA-4T and LEC-4T tumors (**Supplementary Fig. 6**). The reasons for this are unknown. However, numerous studies have demonstrated that tumor cell lines derived from patients often exhibit distinct genetic mutations and characteristics that are not representative of primary tumors (30–32). Many cell lines are derived from late-stage tumors and might have undergone extensive genetic and phenotypic selection subjected to culture conditions. This can lead to reduced immunogenicity because of mutations that do not accurately represent the mutational landscape found in patient tumors (33, 34) Consequently, this limits their potential for translation and the ability to replicate the immune responses seen *in vivo*.

Intriguingly, soluble FasL was only found in the CM of RAS-induced senescent HDF. The reason for this may be that the release of soluble FasL requires the action of metalloproteinases (MMP) (29) and that RAS-induced senescent HDF expresses much higher MMP levels than IR-induced senescent HDF (3). As MMP can also be highly expressed by cancer cells (35–37), it is possible that soluble FasL is released from all types of senescent cells in the TME. Soluble FasL has been reported to inhibit Fas-mediated apoptosis induced by cytotoxic T cells, suggesting that senescent HDF can protect tumor cells by preventing their Fas-mediated killing by T cells or by inducing apoptosis of immune cells directly (38). Indeed, FasL has emerged as an important factor in the TME that can limit the anti-tumor immune response and the efficacy of immunotherapy by inducing the apoptosis of T cells (39, 40). Consistent with these results, we found that ablating FasL expression on senescent HDF abrogated their ability to induce T and NK cells apoptosis and consequently enhanced tumor cell survival *in vitro* and tumor rejection in mice. Interestingly, Motz et al. reported that FasL expression on the tumor endothelium could protect tumor cells (41). Overall, our data support a mechanism of tumor resistance where FasL expressed by senescent stromal cells induces apoptosis of T and NK cells. Given the relatively limited number of immune cells injected in mice and the fact that most NK cells do not survive more than a few days in NSG mice in the absence of appropriate cytokines (for example, human IL-15), it is likely that tumor cells were mostly killed by T cells in our experimental setting. However, we cannot rule out the implication of NK cells, given we previously showed that the adoptive transfer of a limited number of NK cells in NSG mice is sufficient to reject iPSC-derived teratomas (42).

Other immune resistance mechanisms have been reported to be induced by senescence. For example, in mouse syngeneic models, senescent stromal cells can induce an immunosuppressive environment by favoring the formation of myeloid-derived suppressive cells (8). In our study, it was not possible to verify if this mechanism of resistance is also induced in the tumors, given the short half-life of human granulocytes in humanized mice. Another limitation of our study is that it remains to be determined if patient-derived tumors upregulate the expression of FasL in response to therapy-induced senescence, as detected in our xenotransplanted mice. Moreover, more experiments will be necessary to understand the extent and in which context the increased expression of FasL occurs. Indeed, we found that FasL expression is not regulated at the transcriptional level during senescence, and its upregulation was not observed in publicly available RNAseq datasets. The relationship between the molecular pathways involved in senescence and the expression of FasL also remains to be elucidated. We speculate that cytotoxic stress and DNA-damaging agents that can trigger the activation of the c-Jun N-terminal kinase (JNK) pathway may be involved since this pathway was shown to regulate FasL expression (43).

In summary, our results uncover a role for FasL during senescence and suggest it plays an important role in the tumor immune response. It will be interesting to determine the extent to which FasL expression is increased in the context of therapy-induced senescence in different types of human tumors and if targeting FasL can improve the efficiency of immunotherapy.

## METHODS

### Cell culture

Human fibroblasts cell lines (BJ, WI-38 and IMR-90) were a gift from Dr. Judith Campisi or derived from the skin of a healthy adult male donor (41 years old) as previously described (25, 44). Adult skin biopsy was collected in accordance with the ethics committee from the Centre Hospitalier Universitaire (CHU) Sainte-Justine (protocol 2017-1476). The biopsy was cleaned out, cut into 1–5-mm^2^ pieces, and digested with collagenase D (Roche) for 1 h at 37°C with agitation, centrifugated at 400 × *g* for 5 min, and washed with DMEM (Wisent Bio Products). HDF were maintained in DMEM with 10% FBS and 0.2% primocin. HDF senescence was induced by exposing cells to a 12Gy dose of ionizing radiation (1 Gy/min using a Faxitron CP-160), after their stable transduction with lentiviral particles carrying an inducible version of K-RAS, or after treatment with 0.1µM of Doxorubicin. Conditioned media (CM) was prepared from non-senescent or senescent HDF cultured in monolayer with RPMI 1640 (Wisent Bio Products) without serum for 24 hours after which it was then collected, filtered, and centrifuged. HDF were counted at the time of collection, and the CM normalized to the lowest amount of cells in a particular condition by dilution with fresh RPMI.

### iPSC-derived tumor models

iPSCs reprogramming and cellular transformation were performed as previously described (25). In brief, iPSC-derived hepatocyte progenitors (HEPA-4T) and iPSC-derived lung epithelial progenitors (LEC-4T) were generated and transduced using lentiviral particles carrying the SV40 large T antigen and the neomycin resistance gene. Three days later, 300 μg/ml G418 (Thermo Fisher Scientific) selection was applied. Surviving cells were subsequently transduced with lentiviral particles carrying the H-RAS^V12^ and puromycin-resistance genes. Cells were selected for three days post-transduction with 2 μg/mL of puromycin (Thermo Fisher Scientific). Surviving cells were then transduced with lentiviral particles carrying the human telomerase gene. Finally, transformed cells were transduced to express the mPlum gene. All transductions were carried out using fresh culture media containing 8 μg/ml polybrene (Sigma-Aldrich).

### Animals and solid tumor models

Animal experiments were performed under a protocol approved by the institutional committee for good laboratory practices for animal research (protocol #2022-3508). Nine-week-old female and male NSG-SGM3 (expressing human IL3, GM-CSF, and SCF) mice originally obtained from The Jackson Laboratory (Bar Harbor, ME) and bred at the animal care facility at the CHU Sainte-Justine Research Center were used. Mice were housed under strict specific pathogen-free conditions and handled with aseptic techniques under anesthesia (2% isoflurane) to inject tumor cells.

For the subcutaneous injections, 5 × 10^4^ transformed cells (HEPA-4T or LEC-4T) alone or mixed with 2 × 10^5^ non-senescent or senescent HDF were diluted in 100 μl of RPMI 1640 (Wisent Bio Products) and implanted in the right and left flank of mice previously anesthetized and shaved. Tumor growth was monitored for four to five weeks using the Q-Lumi In Vivo imaging system (MediLumine, Montreal, QC, Canada) by fluorescent tracking of mPlum-expressing tumor cells (Ex. 562-40 nm and Em. 641-75 nm). Non-senescent or senescent HDF were stained with a fluorescent dye before their injection in mice. In brief, cells were diluted at 1 × 10^6^ cells/ml, and CellBrite NIR790 Cytoplasmic Membrane Dye (catalog 30079; Biotium) was added at the concentration of 1 μM and incubated at 37°C for 20 min. Cells were then washed twice in serum-free RPMI 1640 (Wisent Bio Products) before being resuspended in 100µL of cold serum-free RPMI 1640 (Wisent Bio Products) for injection in mice. HDF were tracked using near-infrared filters (excitation 769-41 nm and emission 832-37 nm). The tumor fluorescent signal was analyzed and normalized using Fiji macros for picture processing and expressed in fluorescence-integrated density or radiance (photons · s^−1^ · sr^−1^ · cm^−2^) integrated density.

For tumor growth analysis, mice were sacrificed when one limit point was reached according to our animal comity guidelines. Our comity established limit points as no more than 10% weight loss, no distress signs such as alopecia or decreasing activity and that tumor size does not reach more than 1500 mm^3^ or become ulcerated. Tumors were resected and weighed after sacrifice. Tumors were considered eliminated when they were unpalpable or too small to be harvested at sacrifice.

### Targeted irradiation in mice

For whole-thorax irradiation treatments, 29 days after the intravenous injection of 4×10^5^ non-senescent HDF, mice were anesthetized with a mixture of ketamine (100 mg/kg) and xylazine (10 mg/kg), then placed in a lead-shielded container exposing only the thoracic region. A single dose of 12 Gy X-rays was administered. Seven days later, mice were euthanized and their lungs were harvested and frozen for analysis of FasL expression on the injected HDF. Frozen lungs were embedded in OCT compound, and 12 µm sections were prepared using a Leica cryostat. Sections were mounted on gelatin-coated slides and subjected to immunofluorescence staining using a GFP polyclonal antibody (A-11122; Thermo Fisher Scientific, USA), a FasL antibody (NOK-1, sc-33716; Santa Cruz Biotechnology), and DAPI for nuclear counterstaining. Tissue sections were imaged using a Zeiss LSM710 confocal microscope with a 63× objective

### Mouse immune reconstitution and characterization of the tumor-immune infiltrate

For the adoptive transfer of human immune cells, peripheral blood mononuclear cells (PBMCs) were isolated using the Ficoll-Paque gradient (GE Healthcare) from the blood of healthy donors after informed consent. Granulocytes were collected by lysing the red blood cells (RBC) by resuspending the Ficoll-Paque cell pellet in 19mL of sterile deionized water for 20 seconds before adding 1 ml of sterile 20× PBS solution. PBMCs and granulocytes were counted and mixed at a 1:1 ratio before being injected into mice; a total of 1 × 10^7^ human immune cells (5 × 10^6^ each) were injected i.p. with 200 μl of RPMI 1640 (Wisent Bio Products). Tumors were digested using the human Tumor Dissociation Kit and gentleMACS Octo Dissociation with Heaters (Miltenyi Biotec), according to the manufacturer’s instructions. The cell suspension was filtered on 40 μm MACS SmartStrainers (Miltenyi Biotec) and washed using RPMI 1640 (Wisent Bio Products) containing 10% FBS. Finally, resuspended cells were stained with the following antibodies to analyze the tumor-infiltrating immune cell: mouse CD45/PE/Cy7, human CD45/BUV395, human CD3/AF700, human CD19/PE-CF594, human CD4/BB515, human CD8/BV421, human CD14/APC/H7, human CD56/BV786, and human CD127/BB700, all from BD Biosciences except human CD33/BV510 and human CD25/BV711 who were purchased from Biolegend. All data were acquired on LSR Fortessa (BD Biosciences) and data analysis was done on FlowJo V10 (v10; Tree Star).

### CRISPR-Cas9 knockout cell lines

FasL Knock-out (KO) HDF and A549 cells were generated by the Gene Editing Platform of the CHU Sainte-Justine Research Center using CRISPR-Cas-mediated genome editing. Briefly, a guide RNA (gRNA) targeting a DNA sequence within the first exon of the FasL gene (5’-CTGGGCACAGAGGTTGGACA-3’) was cloned into the BsmB*I* restriction site of the 3^rd^ generation LentiCRISPR.V2 (Addgene, #52961) vector. Lentiviral particles were produced by transfecting LentiCRISPR.V2 (pLC-EF1a-SpCas9-U6-gRNA1-FasL 450 ng) together with the pMDL (750 ng), pREV (300 ng), and pVSV-G (390 ng) packaging plasmids in 5 × 10^5^ HEK293T cells with Lipofectamine 2000 (Invitrogen). The supernatant was collected 40 hours later, centrifuged 5min at 3000 rpm on a Sorvall LegendMicro17 (ThermoFisher) to remove debris and then frozen at - 80°C. FasL KO HDF were produced by transducing 5 × 10^5^ cells with 100 µL of viral particles. After 24 hours, Puromycin (2 mg/mL) (Wisent) was added to select positive clones expressing Cas9 and the gRNA. 72 hours later, 10 cells were transferred into separate wells of a 96-well plate and left to establish a positive clone mixture. PCR and sequencing were performed to characterize knock-out populations. Finally, ICE Analysis Software (Synthego) showed that 1 cell population no longer expressed FasL. The latter was composed of 37% and 63% of cells with indels of +1 bp and -11 bp, respectively. The loss of expression of FasL was confirmed by FACS analysis (LSR Fortessa) (BD Biosciences) using a FasL/PE conjugated antibody from BioLegend.

### 3D spheroids formation, invasion assay, and image analysis

Spheroids were formed by centrifugating 5000 tumor cells/well (LEC-4T, HEPA-4T, A549) alone or together with 5000 HDF/well (non-senescent and senescent) for 5 min at 150g in ultra-low attachment 96-well plates and incubated at 37°C under 5% CO^2^ for 6 days. Where indicated, 5 × 10^5^ peripheral blood mononuclear cells (PBMCs) were added per well, and 24 or 48 hours later, spheroids were dissociated into single cells for subsequent flow cytometric analysis. Alternatively, spheroids were embedded in OCT, frozen and 12 µm sections were generated using a Leica cryostat. Sections were transferred to gelatinized slides and immunofluorescence staining was performed using a human CD45 antibody (Rat. YAML501.4; Thermo Fisher Scientific, USA), GFP Polyclonal Antibody (A-11122; Thermo Fisher Scientific, USA), FAS-L Antibody (NOK-1. sc-33716; Santa Cruz Biotechnology), and DAPI to counterstain DNA.

### 3D tumor spheroids growth after therapy-induced senescence

Spheroids (A549 mPlum cells alone or together with GFP-expressing HDF) were allowed to form for five days after which they were treated with 0.1µM of doxorubicin or a single dose of 15 Gy IR. Spheroids size was monitored using the IncuCyte S3 (Essen Bioscience) through 4X images taken every 3 hours for a total of 96 hours. Likewise, some spheroids were fixed with 4% formaldehyde and stained for SA-β-gal activity.

### Immunophenotyping of HDF by Flow Cytometry

HDF were harvested, re-suspended in phosphate-buffered saline (PBS, Wisent Bio Products), and stained with the LIVE/DEAD™ Fixable Far-Red Dead Cell Stain Kit (Thermo Fisher Scientific) for 15 min on ice. Cells were then washed twice and re-suspended in flow cytometry staining buffer (BD Biosciences) and incubated for 30 min at 4°C with 1 µL of PE anti-human FasL antibody (clone NOK-1) from BioLegend. Nonspecific background signals were measured by incubating cells with the appropriate isotype-matched antibody. Unstained, viability dye only, and single-stained compensation beads (Invitrogen) served as controls. Doublets were gated out using forward-scatter width/height and sideward-scatter width/height event characteristics. All stained cells were analyzed using an LSR Fortessa cytometer (BD Bioscience), and the obtained results were analyzed using FlowJo v10 (v10; Tree Star).

### Detection of apoptosis

To detect apoptotic cells we used the IncuCyte S3 (Essen Bioscience) live cell imaging system in combination with propidium iodide (PI) or the caspase 3/7 specific activity dye. In brief, PBMCs (3 × 10^5^) were co-culture with non-senescent or senescent HDF (3 × 10^4^) in a 48-well plate. Alternatively, PBMCs were cultured in the presence of CM collected from HDF in a 48-well plate. PI (25µg) and IL-2 (150 IU/mL) were added to the media and cell viability was monitored for 72 hours. Cells were treated with 7 µM of camptothecin as a positive control of cell death. Dead cells were counted via red object counts at indicated time points, and data was analyzed using the IncuCyte software. To measure apoptosis of tumor cells we used the caspase 3/7 activity dye. Tumor cells (1 × 10^4^) were seeded in a 24-well plate and treated with 200 ng/mL IFN-γ overnight. The next day, PBMCs were added at a ratio of 1:20 (tumor cells: PBMCs) in the presence or absence of CM collected from non-senescent or senescent HDF and the Incucyte® green Caspase-3/7 activity reagent (Sartorius, Ann Arbor, MI, USA). Plates were analyzed at a two-hour interval for a total of 72 hours.

The apoptosis of PBMCs was also measured by flow cytometry using the Apotracker(TM) Green (Biolegend) apoptosis probe in combination with PI (Sigma Aldrich). Both floating and adherent cells were collected after 3 days of co-culture, washed twice with PBS, and stained with Apotracker(TM) Green following the manufacturer’s protocol. Cells were stained with the following antibodies: human CD45/BUV395, human CD3/AF700, human CD8/BV421, and human CD56/BV786 from BioLegend and then immediately analyzed using the LSR Fortessa (BD Biosciences). Cells were considered alive when not stained with PI or Apotracker, while apoptotic cells were defined as those stained with Apotracker. Absolute counting beads (Invitrogen) were used to normalize FACS event acquisition and to calculate absolute cell numbers.

### NK cell culture conditions and cytotoxicity assay

NK cells were isolated from PBMCs of healthy donors by depleting CD3^+^ cells using an EasySep human CD3 positive selection kit II (StemCell Technologies). The negative cell fraction was expanded using irradiated (100 Gy) K562 mbIL21 4-1BBL for a minimum of 2 weeks in RPMI1640 supplemented with 10% FBS, 1% penicillin/streptomycin (Wisent Bio Products), and 100 UI/mL IL-2 (SteriMax) until the proportion of NK cells (CD3^−^, CD56^+^) reached close to 100% as determined by flow cytometry. Cells were rested for one week after their expansion prior to being used in a cytotoxicity assay. In brief, A549 mPlum cells were co-cultured with GFP-expressing HDF (non-senescent or senescent) at a 1:2 ratio in a 96 well plate for 72 hours. NK cells were then added at different effector: target ratios in the presence of 20 UI/mL IL-2. Cell death were determined by counting cells by flow cytometry. Before the acquisition, the supernatant was gently removed, and cells were stained with Zombie NIR™ Fixable Viability Kit (BioLegend) according to the manufacturer’s instructions and then detached using trypsin. Cytotoxicity was assessed by flow cytometry and was calculated as Lysis (%) = [1-live targets (sample)/live targets (control)] × 100%. Cytotoxicity against A549 was performed using NK cells isolated from the same donor in the presence of non-senescent and senescent HDF. A total of three different donors of NK cells were used in total.

### Statistical analysis

Statistical analyses were performed using GraphPad Prism 8.0 (GraphPad Software, Inc., San Diego, CA). Data are represented as mean ± SEM. For comparisons of three or more groups, data were subjected to t-tests or one-way ANOVA analysis, followed by Dunnett’s multiple comparisons test when comparing every mean to a control mean or Tukey’s multiple comparisons test when comparing every mean to every other mean. The tumor growth kinetics was analyzed using mixed-effects, followed by Tukey’s multiple comparison test. *p < 0.05, **p < 0.01, *** p < 0.001, and ****p < 0.0001.

## AUTHOR CONTRIBUTIONS

M.C., J. D., G.M.N., A.S., performed the experiments; G.M.B., and B.B., provided study material; O.L., provided technical guidance; C.B., designed and supervised the study; M.C., and C.B., wrote the manuscript with the contributions from all the authors.

