## Supplemental Figures 1-6 for "Senescent human fibroblasts have increased FasL expression and impair the tumor immune response"

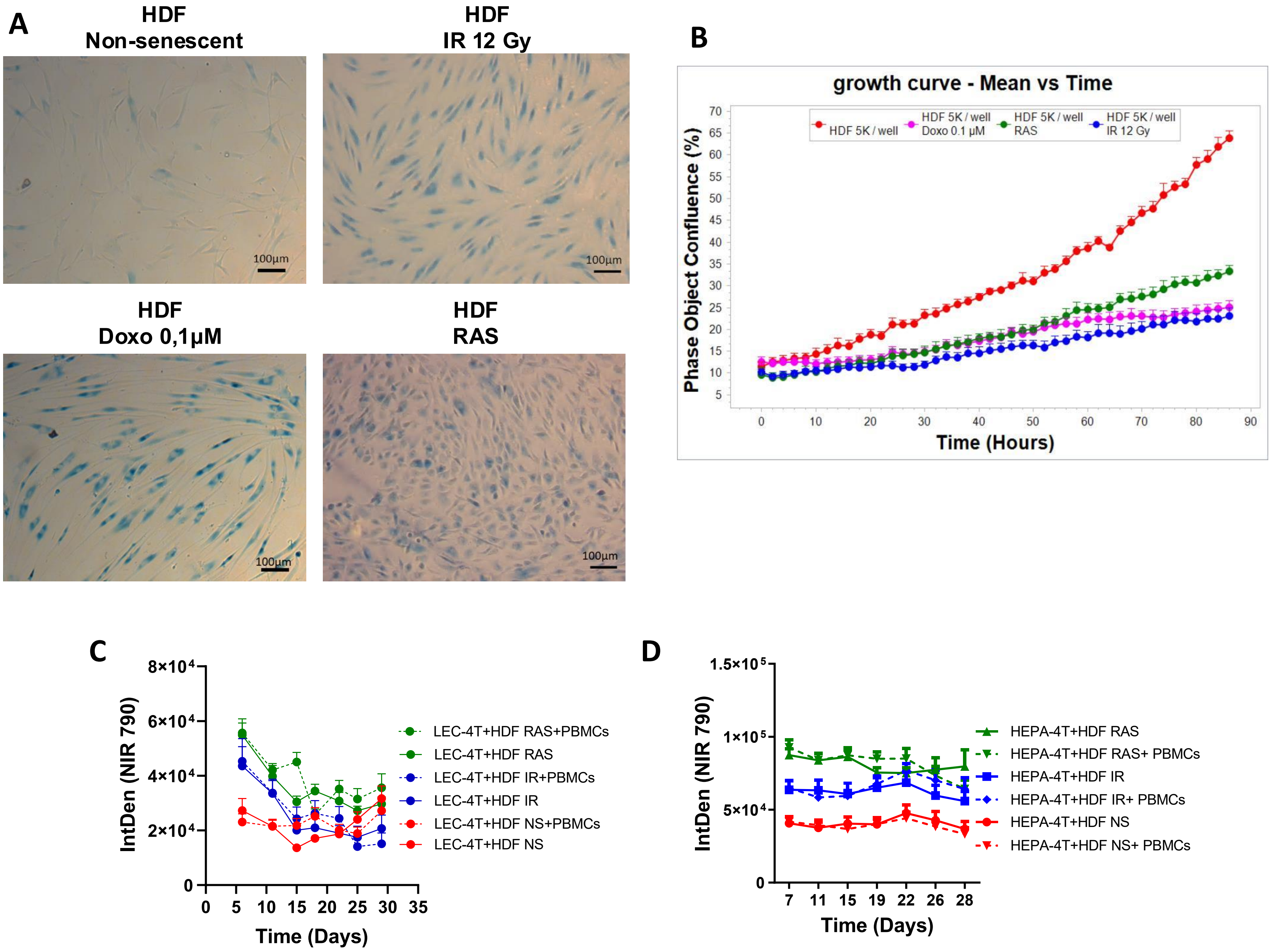


**Supplementary Fig. 1. HDF undergoes senescence *in vitro* and are not eliminated *in vivo*. A)** Representative images of β-galactosidase activity detected on HDF either non-senescent or 7 days after exposure to 12 Gy of IR, 0.1µM of doxorubicin, or expression of RAS. The scale bar represents 100 µm**. B)** Growth curves of non-senescent HDF (red line) and HDF after exposure to IR (blue line), doxorubicin (pink line), and expression of RAS (green line). Shown is the mean ± SEM**. C and D)** Graphs representing the integrated density of HDF stained with NIR790 and tracked over time in LEC-4T or HEPA-4T tumors growing in NSG-SGM3 mice injected or not with autologous immune cells (from tumors in Fig. 1C and D). Non-senescent (NS) HDF are shown in red, HDF exposed to IR in blue and expressing RAS in green. Shown is the mean ± SEM of counts from 12-20 tumors.

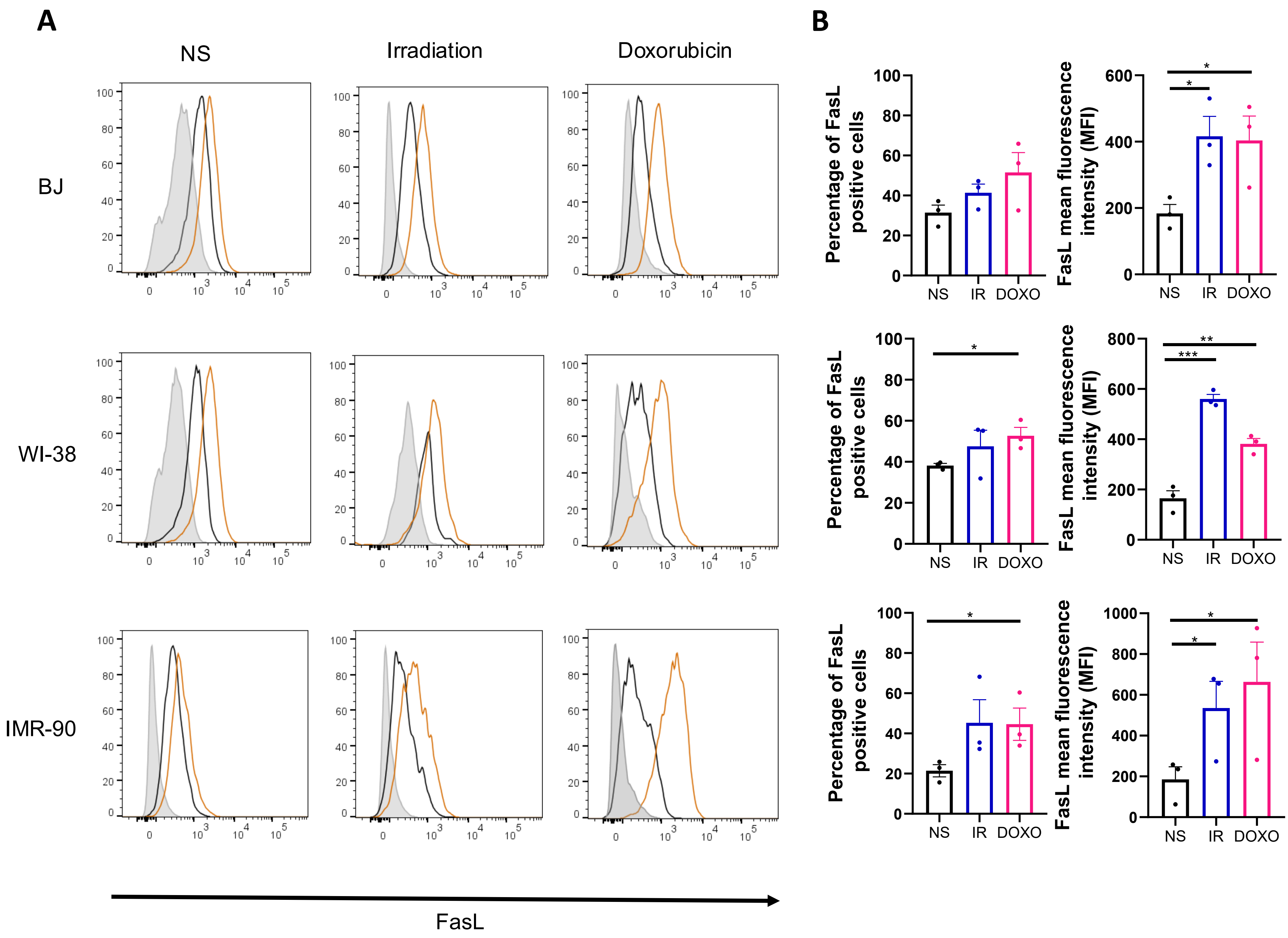


**Supplementary Fig. 2. Increased expression of FasL in different senescent human fibroblast cell lines. A)** Representative histograms showing the expression of FasL as detected by flow cytometry on non-senescent (NS) or senescent (exposed to IR or doxorubicin) on BJ (skin fibroblast), WI-38, and IMR-90 (lung fibroblasts). Also shown are non-stained cells (in gray) and cells stained with the isotype controls (in black). **B)** Bar graphs showing the proportion and the mean fluorescent intensity (MFI) of HDF expressing FasL in the different HDF populations presented in panel A. Shown is the mean ± SEM of three independent experiments. Statistical analysis between groups was performed with one-way ANOVA with Dunnett's multiple comparisons tests.

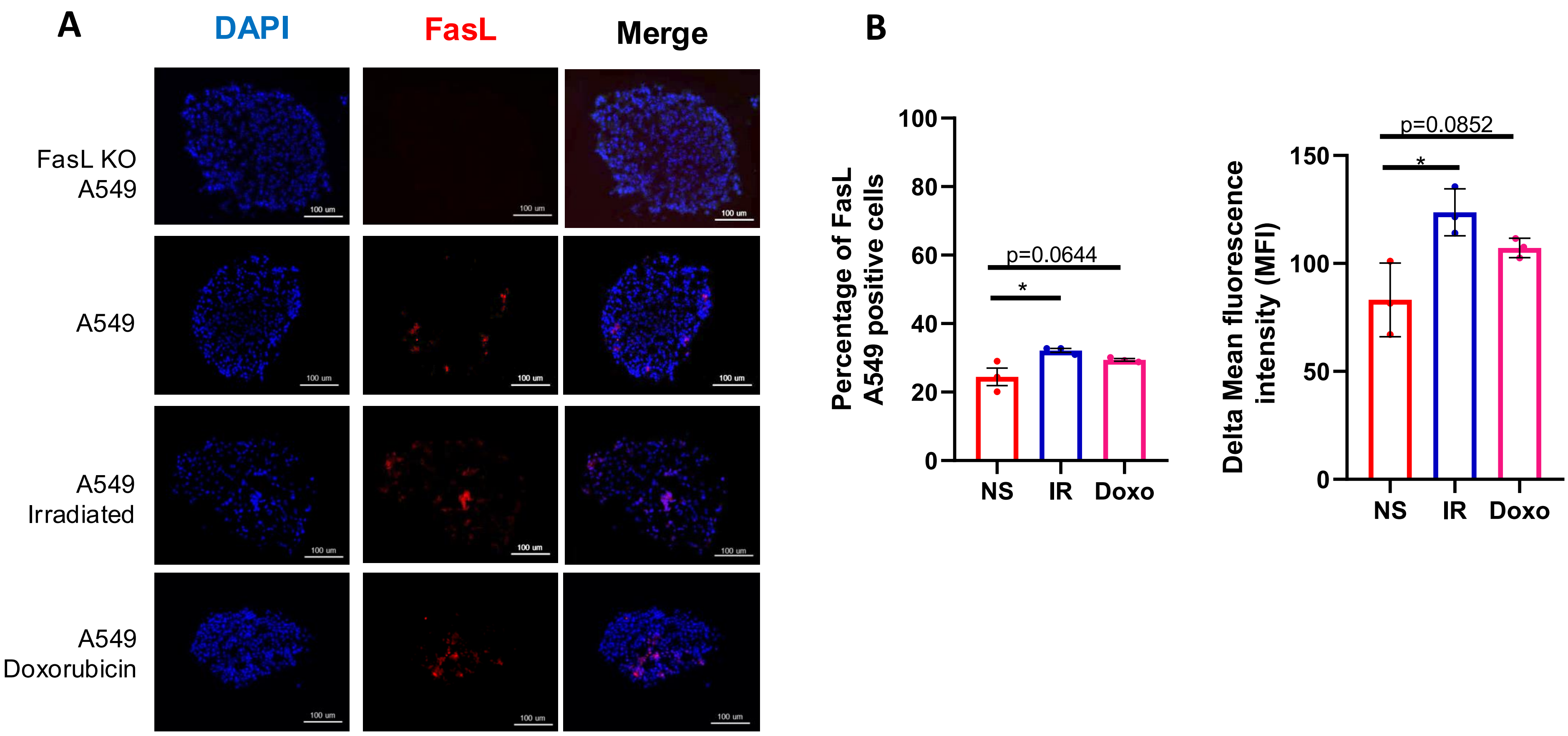


**Supplementary Fig. 3. FasL expression induced in senescent A549 spheroids**. **A)** Representative images of FasL expression (in red) as detected by immunostaining on sections of A549 spheroids exposed or not to therapy-induced senescence 6 days before. Cell nuclei were stained with DAPI in blue. Spheroids produced using FasL KO A549 tumor cells were used as a negative control. The scale bar represents 100 µm. **B)** Quantification by flow cytometry of the proportion of A549 tumor cells expressing FasL when grown for 48 hours in co-culture with non-senescent or senescent HDF (left panel). Also shown is the mean fluorescence intensity (MFI – right panel). The MFI was calculated by subtracting the fluorescence of the isotype control. Statistical analysis between groups was performed with one-way ANOVA with Dunnett's multiple comparisons tests.

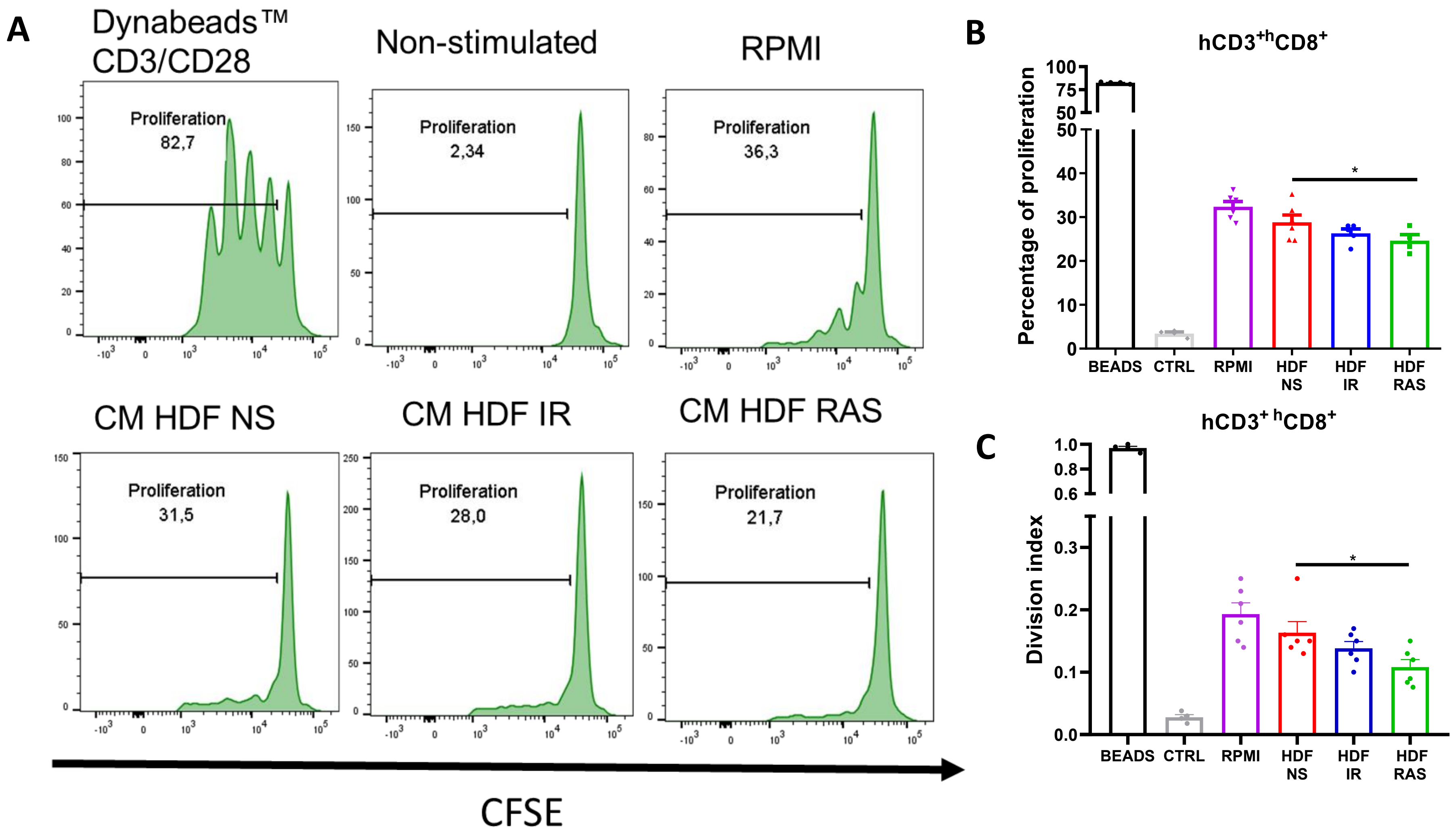


**Supplementary Fig. 4. Proliferation of alloreactive T cells is inhibited by the SASP**. **A)** Representative flow cytometry histograms showing the proliferation of CFSE-labeled responder $CD3^{+}CD8^{+}$ cells in a modified mixed lymphocyte reaction (MLR) assay at a ratio 4:1 (stimulator: responder). Stimulator were irradiated, alloreactive PBMCs and responder were fresh PBMCs. Shown is the proliferation of $CD3^{+}CD8^{+}$ cells after 72 hours when stimulated with CD3/CD28 beads (positive control) when non-stimulated or when stimulated in the presence of RPMI media alone or CM collected from non-senescent or senescent HDF. **B)** Bar graph showing the proportion of cells undergoing at least one cell division from each condition detailed in panel A. **C)** Bar graph showing the division index (mean number of divisions in the total population) of $CD3^{+}CD8^{+}$ cells. Shown is the mean ± SEM of two technical replicates from three independent experiments. Statistical analysis between groups was performed with one-way ANOVA with Dunnett's multiple comparisons test.

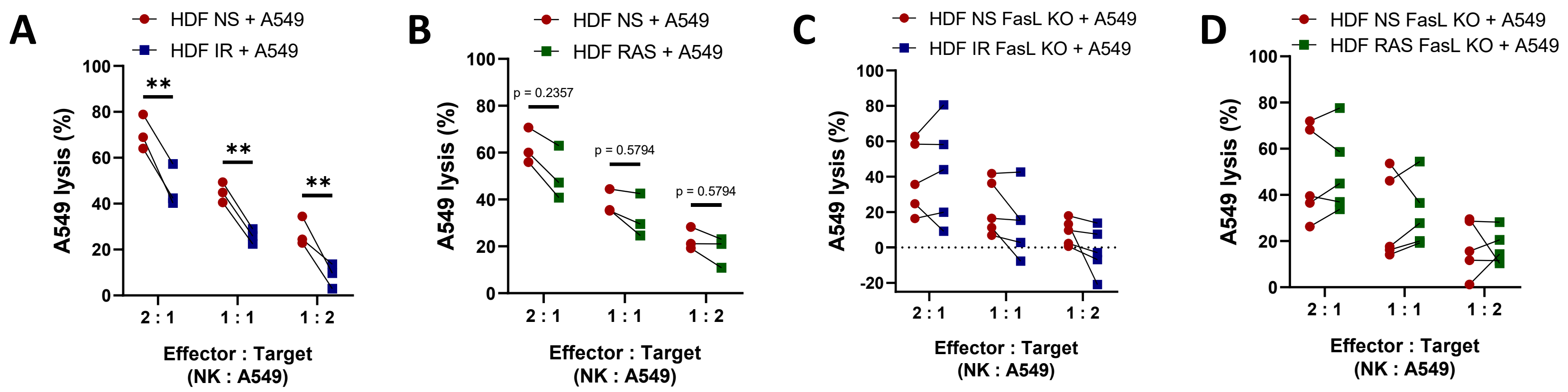


**Supplementary Fig. 5. Senescent HDF protect A549 cells against NK cells**. **A)** NK cells cytotoxicity assay showing the proportion of A549 tumor cells lysis in 24 hours when exposed to purified and activated primary NK cells from 3 distinct donors at different effector to target ratios in the presence of either non-senescent HDF (NS in red) of IR-induced senescent HDF (in blue). Data is presented as paired-matched for each donor. Each dot is the average of two technical replicates. **B**) Same as in panel A, except that A549 lysis was compared to RAS-induced senescent HDF. **C and D)** Same as in panels A and B, except that A549 lysis was evaluated in the presence of FasL KO non-senescent and senescent HDF. Individual values are represented by symbols and connected by a line to show paired data across experimental conditions. The p-value was calculated by multiple paired t-tests.

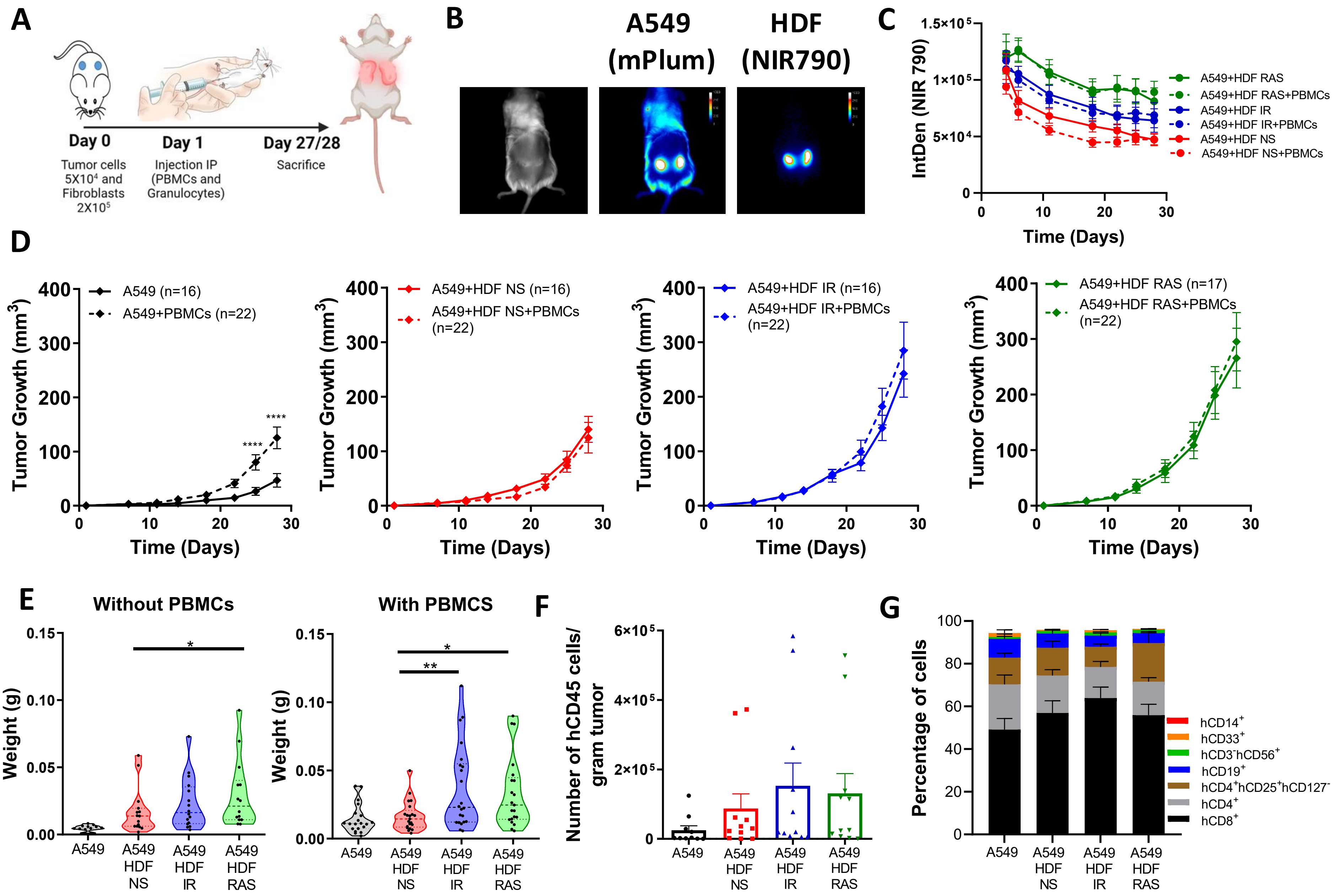


**Supplementary Fig. 6. A549 tumors are resistant to immune clearance.** **A**) Schematic of the experiment. A549 cells (5 × $10^4$ cells) were subcutaneously injected in NSG-SGM3 mice alone or together with non-senescent or senescent HDF (2 x $10^5$ cells) on each flank on day 0. The next day, mice were injected intraperitoneally with PBMCs (5 x $10^6$) and granulocytes (5 x $10^6$). Mice were sacrificed on day 27 or day 28, and tumors were collected for analysis. **B)** Representative images on day 22 of A549 tumor cells expressing mPlum and HDF stained with NIR790 in mice bearing subcutaneous tumors. **C)** Graphs representing the integrated density of HDF stained with NIR790 and tracked over time in LEC-4T or HEPA-4T tumors. **D)** Growth curves for A549 tumors injected alone (in black) or co-injected with non-senescent HDF (in red), senescent HDF induced by irradiation (in blue) or induced by RAS (in green) in mice without (solid line) and with (dashed line) allogenic immune cells. Each line represents the mean tumor growth (± SEM). Statistical analyses were performed using a mixed-effects model, followed by Tukey's multiple comparison test. **E)** Graphs representing the weights of A549 tumors collected at sacrifice. Each dot represents an individual tumor. Values represent the mean ± SEM. A one-way ANOVA with Dunnett's multiple comparisons tests was used to determine statistical significance. **F)** Graph showing the absolute counts of tumor-infiltrating human CD45⁺ cells per gram of dissociated A549 tumors as determined by flow cytometry. Each dot represents the count from a single tumor. Shown is the mean ± SEM. Statistical analysis between groups was performed with one-way ANOVA followed by Dunnett's multiple comparisons tests. **G)** Stacked bar graph showing the proportion of immune subset cell populations in tumors from each group. Shown is the mean ± SEM.
